## supplementary figures for "Strip1 regulates retinal ganglion cell survival by suppressing Jun-mediated apoptosis to promote retinal neural circuit formation"

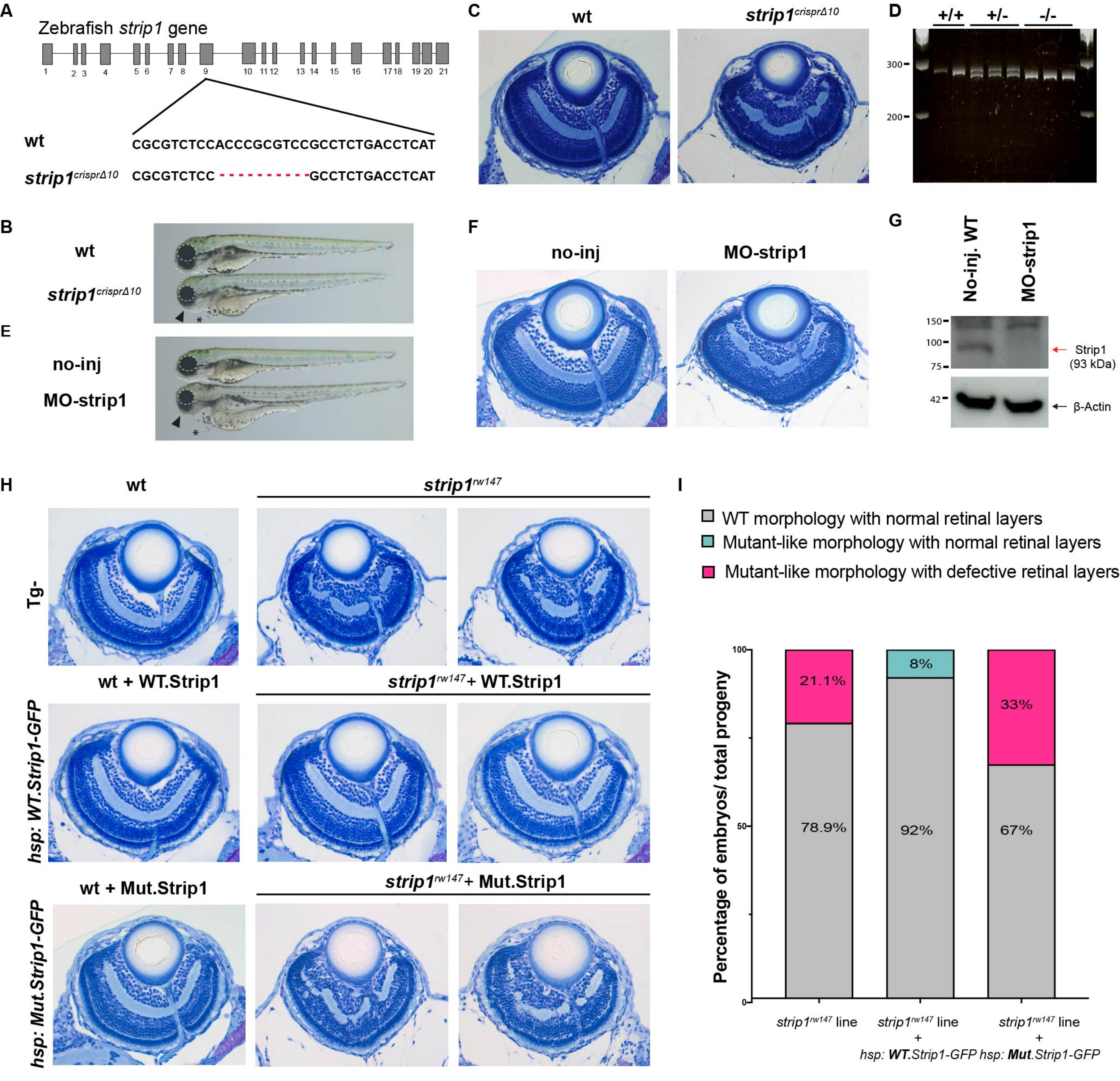


**Figure 1–figure supplement 1.**

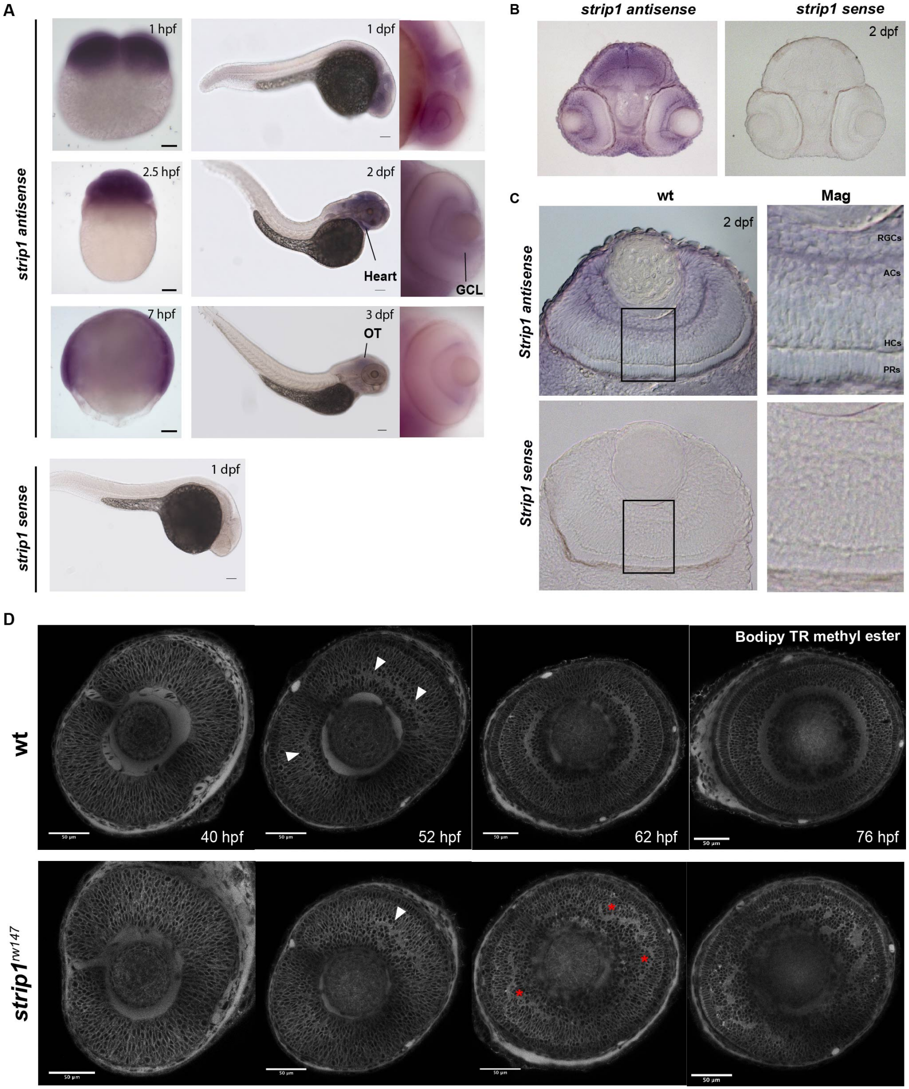


**Figure 1–figure supplement 2.**

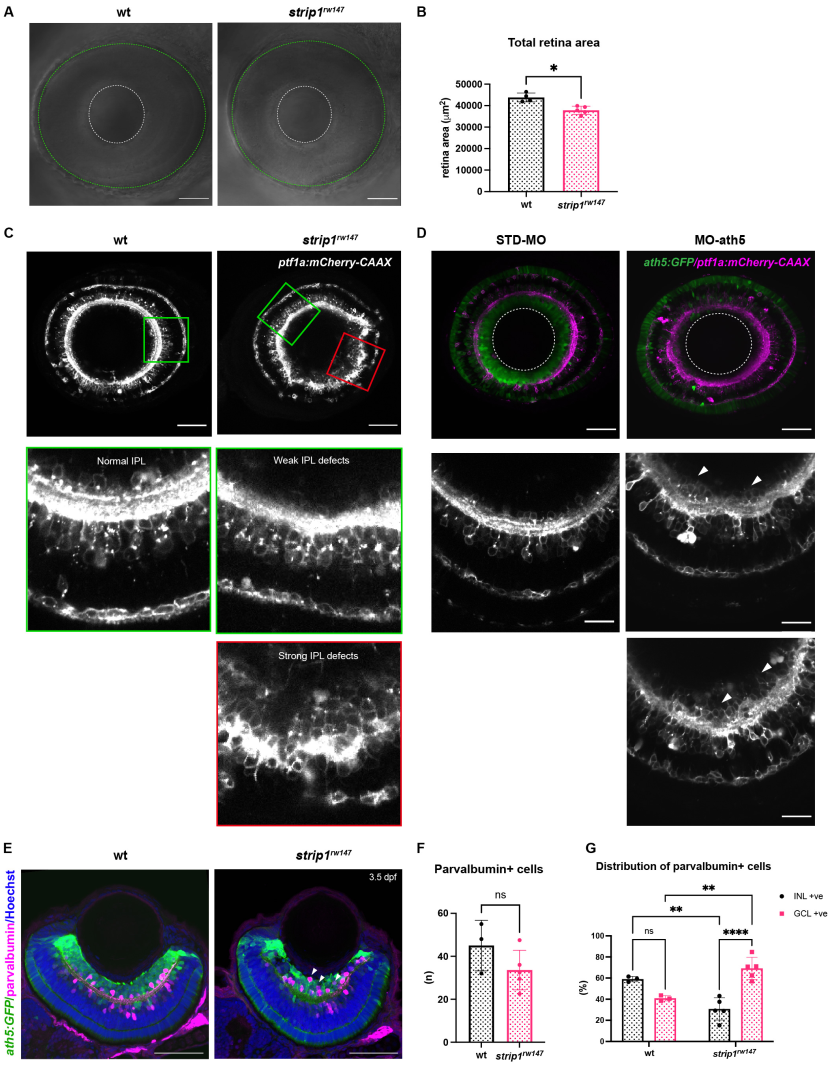


**Figure 2–figure supplement 1**

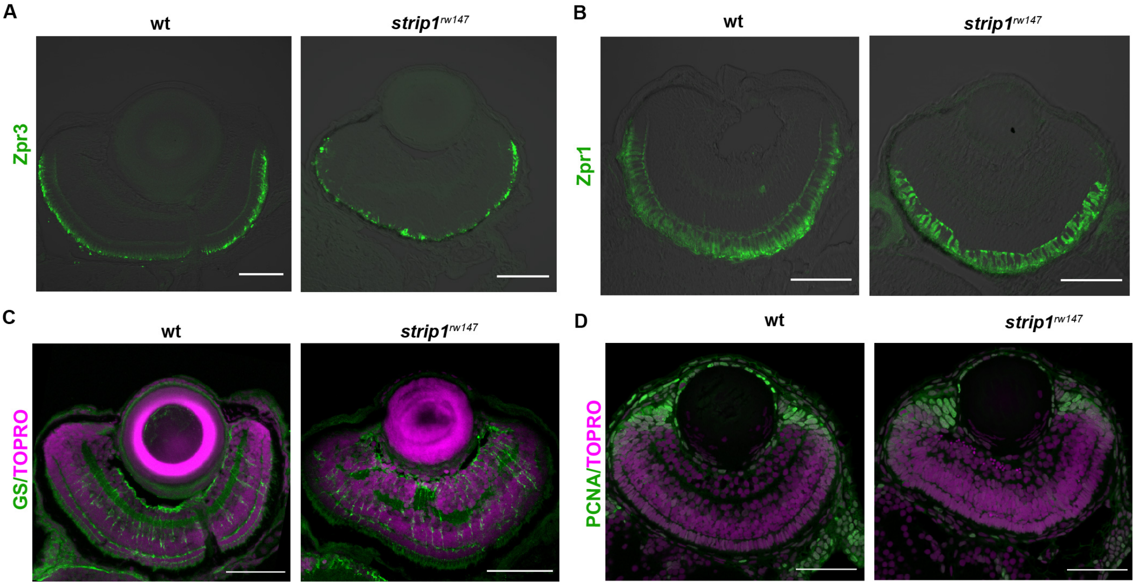


**Figure 2–figure supplement 2**

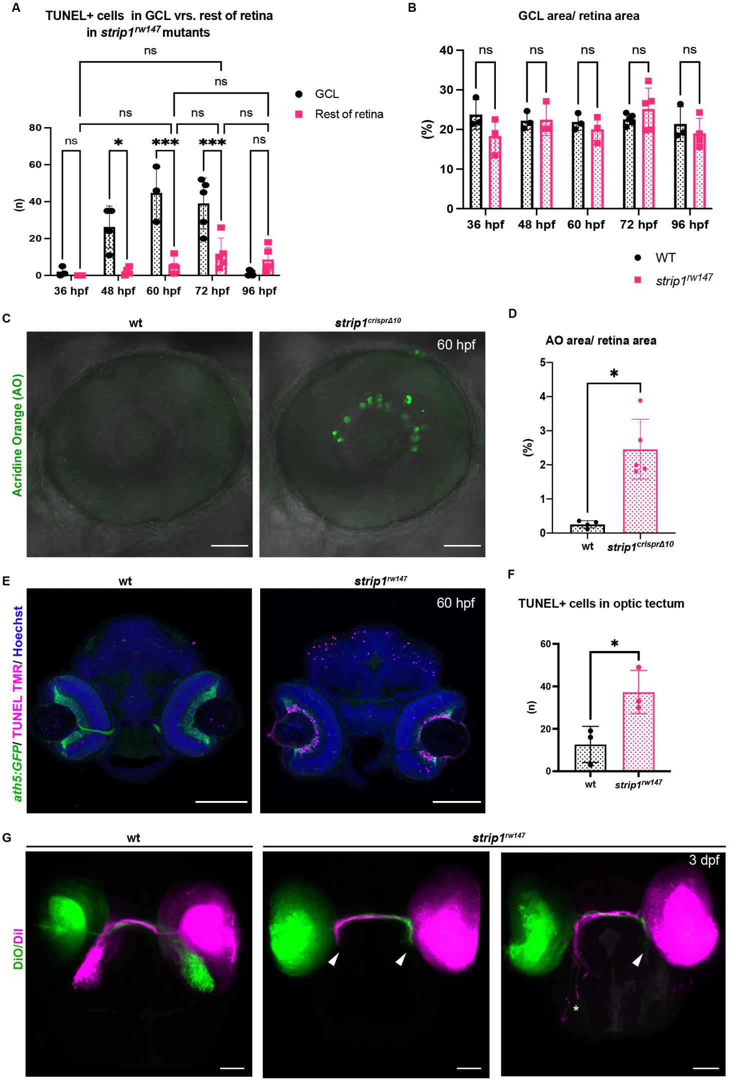


**Figure 3–figure supplement 1.**

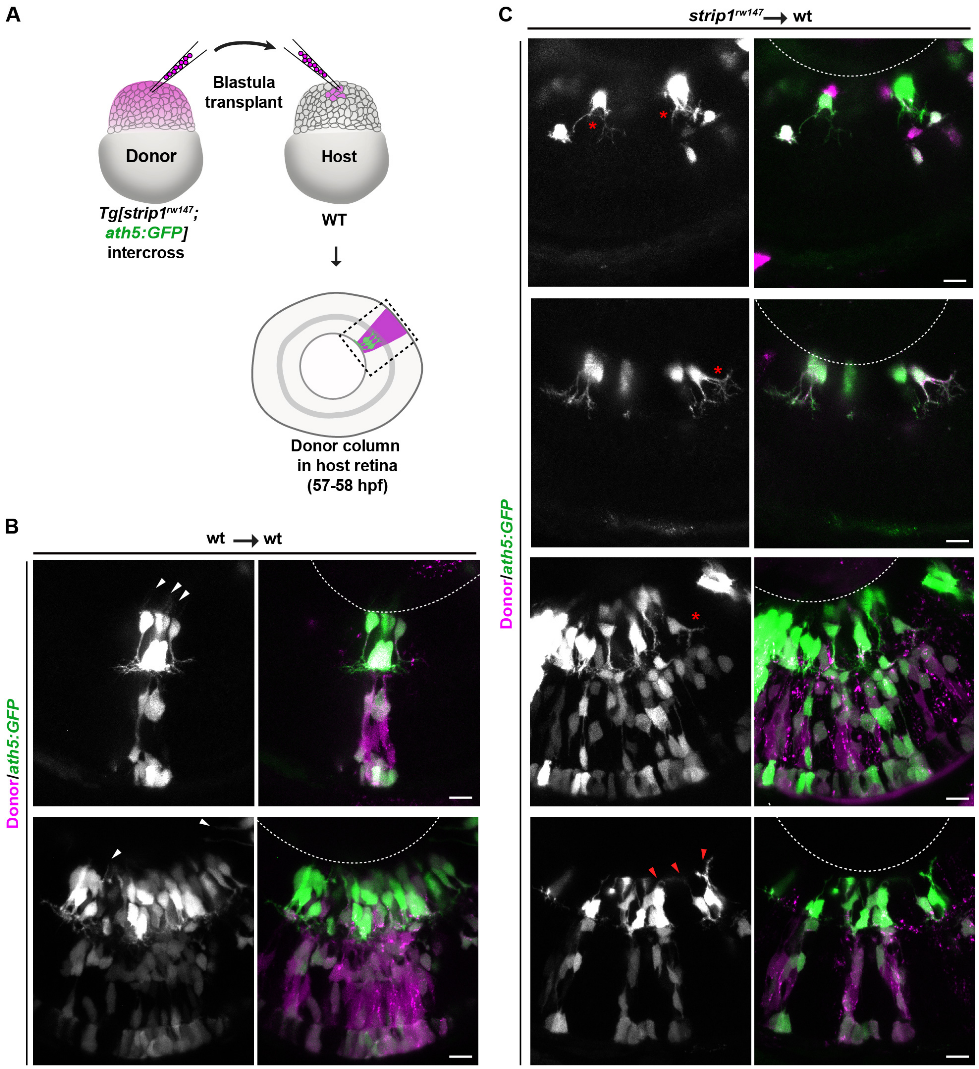


**Figure 3–figure supplement 2**

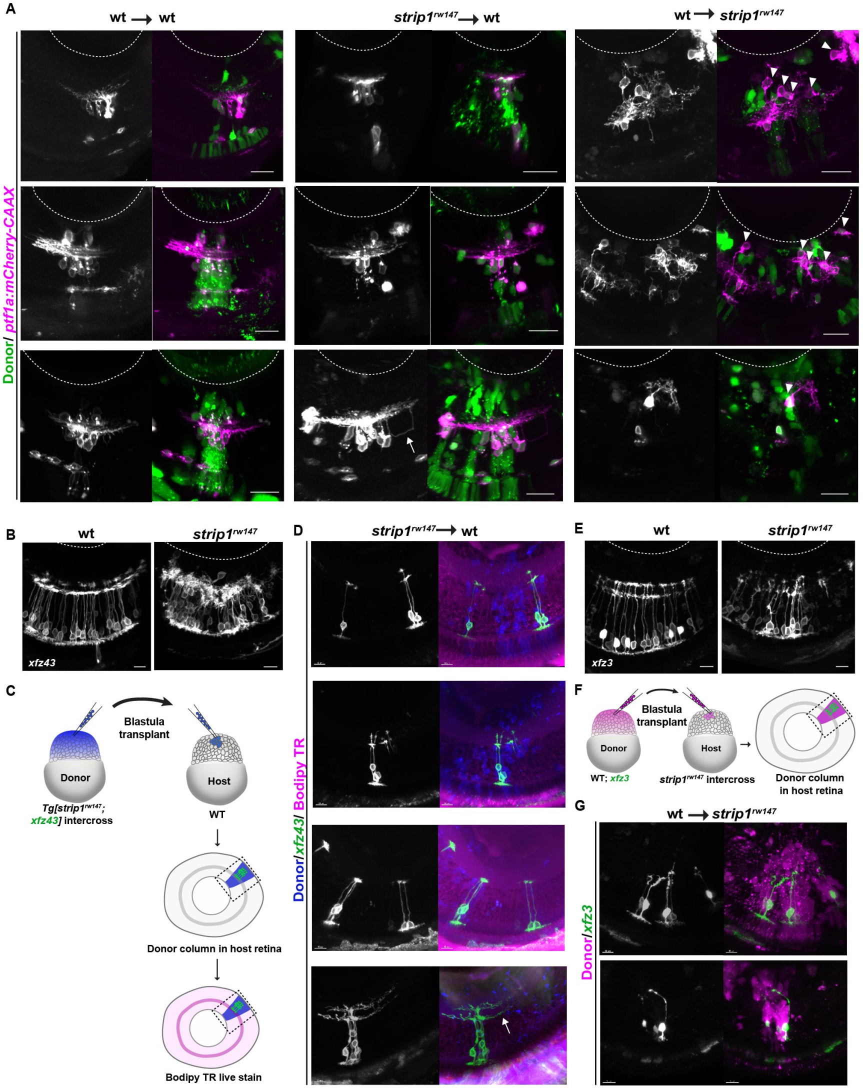


**Figure 4–figure supplement 1.**

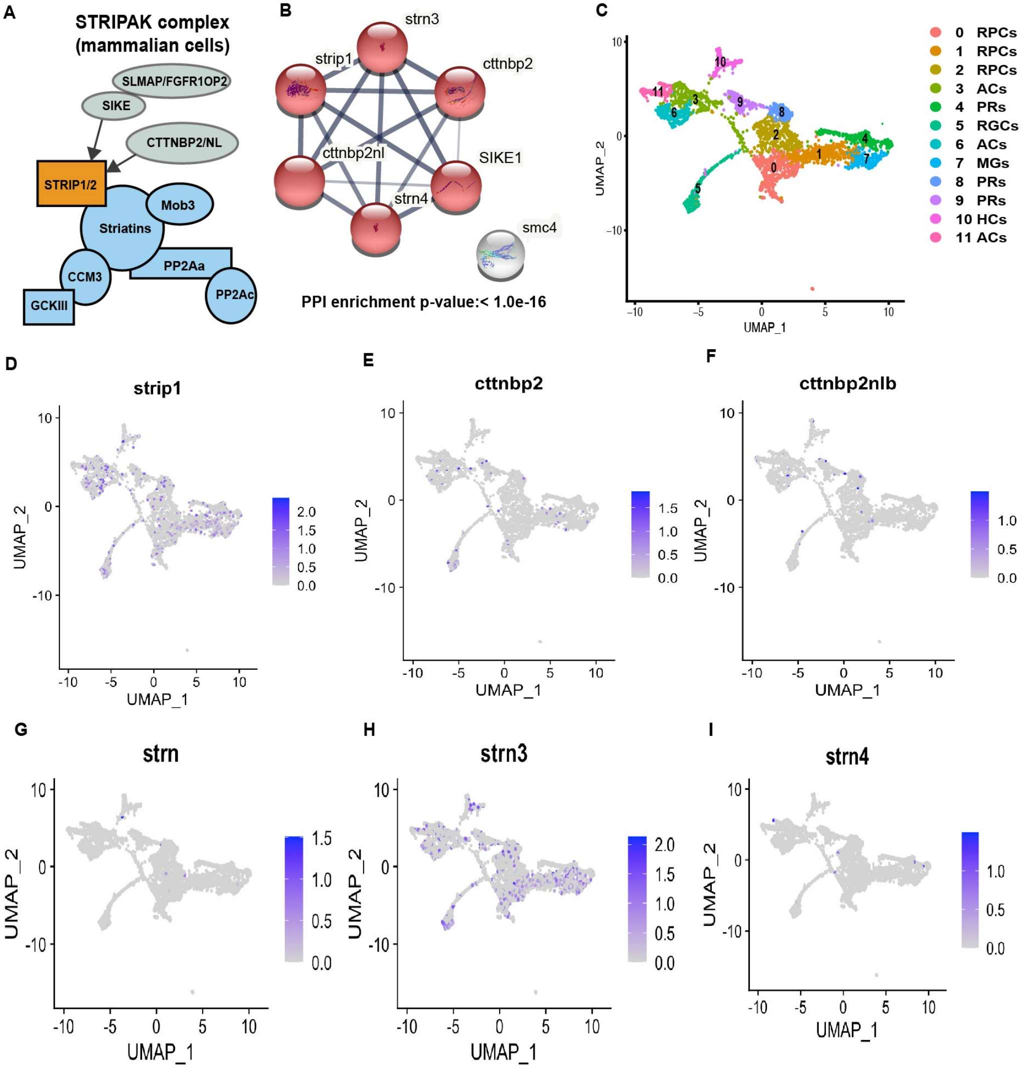


**Figure 5–figure supplement 1.**

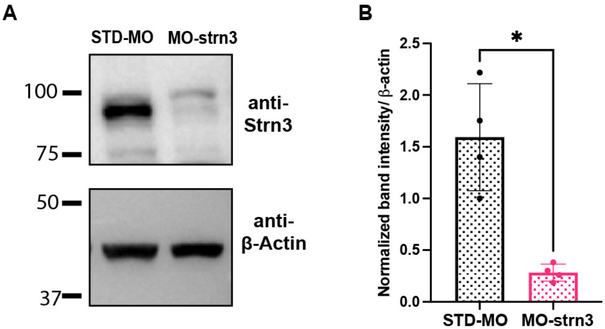


**Figure 5–figure supplement 2**

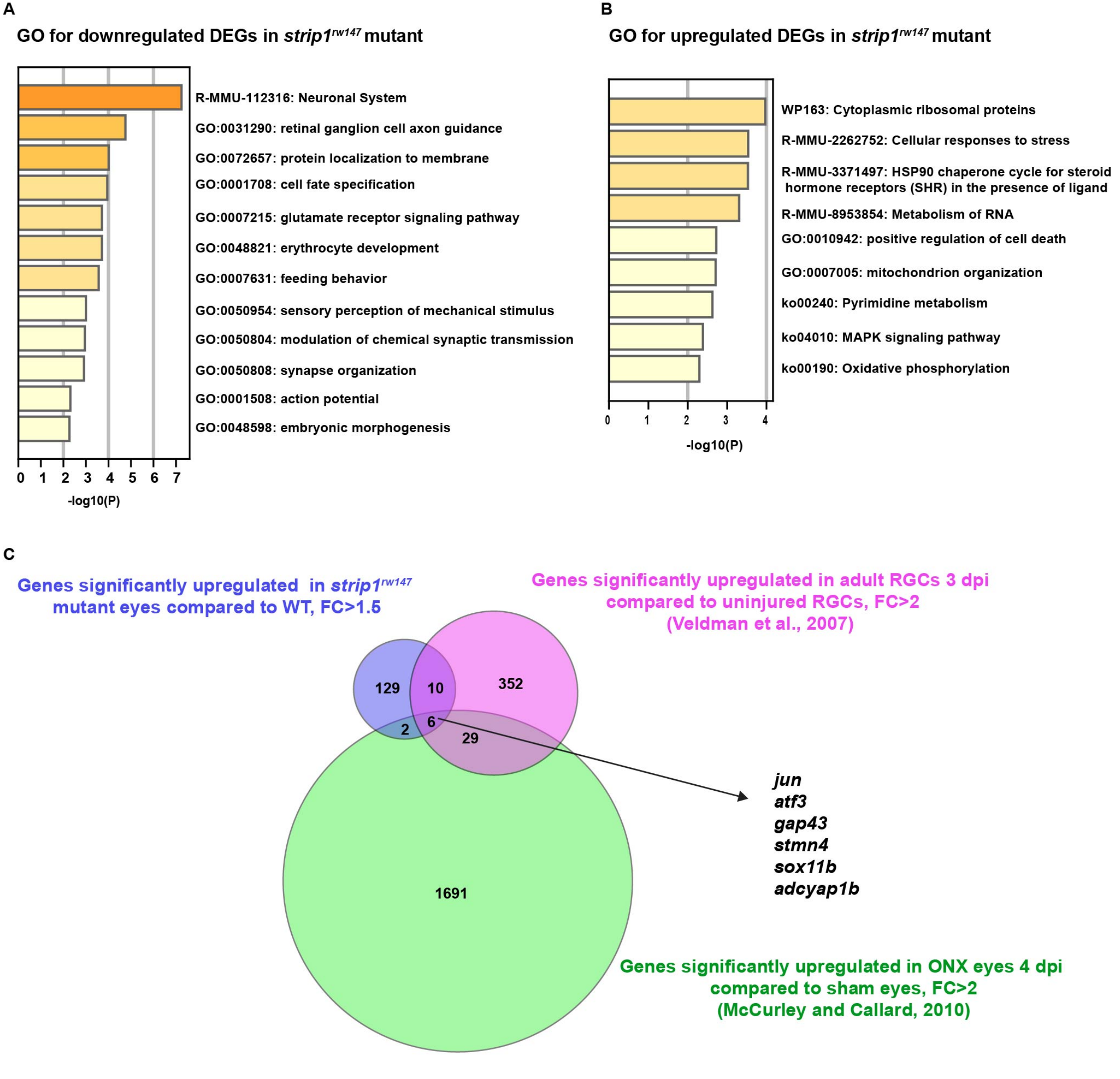


**Figure 6–figure supplement 1.**

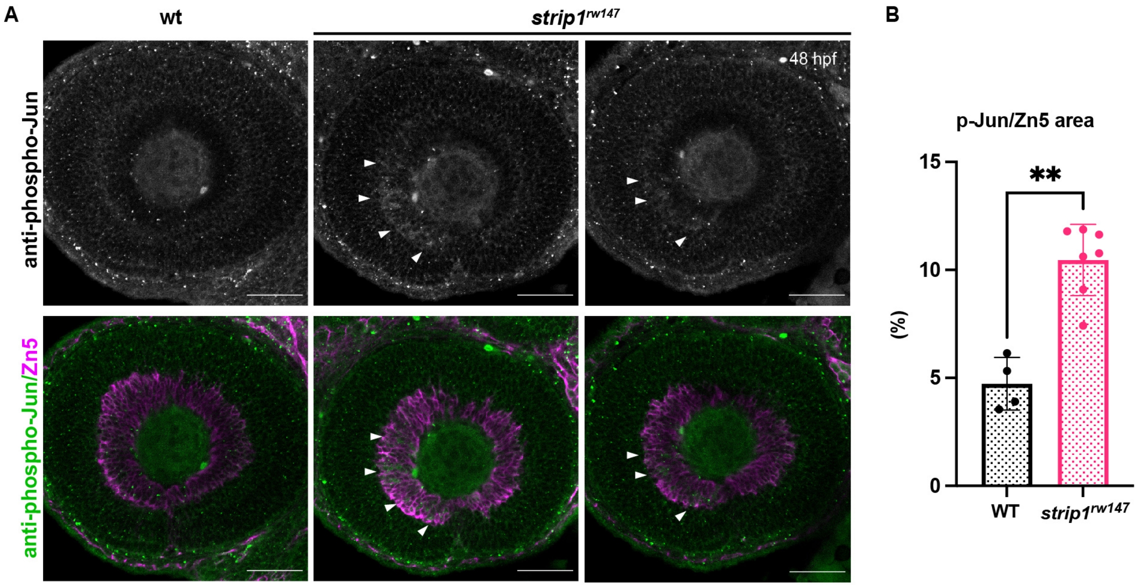


**Figure 6–figure supplement 2**
